# Transgenic mouse lines expressing the 3xFLAG-dCas9 protein for enChIP analysis

**DOI:** 10.1101/221820

**Authors:** Toshitsugu Fujita, Fusako Kitaura, Asami Oji, Naoki Tanigawa, Miyuki Yuno, Masahito Ikawa, Ichiro Taniuchi, Hodaka Fujii

**Author notes:** Corresponding author, Department of Biochemistry and Genome Biology, Hirosaki University Graduate School of Medicine, Zaifu-cho 5, Hirosaki, Aomori, 036-8562 Japan.

## Abstract

We developed the engineered DNA-binding molecule-mediated chromatin immunoprecipitation (enChIP) technology to isolate specific genomic regions while retaining their molecular interactions. In enChIP, the locus of interest is tagged with an engineered DNA-binding molecule, such as a modified form of the clustered regularly interspaced short palindromic repeats (CRISPR) system containing a guide RNA (gRNA) and a catalytically inactive form of Cas9 (dCas9). The locus is then affinity-purified to enable identification of associated molecules. In this study, we generated transgenic mice expressing 3xFLAG-tagged *Streptococcus pyogenes* dCas9 (3xFLAG-dCas9) and retrovirally transduced gRNA into primary CD4^+^ T cells from these mice for enChIP. Using this approach, we achieved high yields of enChIP at the targeted genomic region. Our novel transgenic mouse lines provide a valuable tool for enChIP analysis in primary mouse cells.

## Introduction

Identification of molecules associated with a genomic region of interest *in vivo* is an essential step in understanding the regulatory mechanisms underlying that region’s functions. To this end, we previously developed engineered DNA-binding molecule-mediated chromatin immunoprecipitation (enChIP) technology to isolate genomic regions of interest along with their interacting molecules (Fujita *et al.* 2013; Fujita & Fujii 2013). In enChIP, the locus of interest is tagged with engineered DNA-binding molecules, such as transcription activator-like (TAL) proteins (Bogdanove & Voytas 2011) or a variant of the clustered regularly interspaced short palindromic repeats (CRISPR) system (Harrison *et al.* 2014; Wright *et al.* 2016) containing a guide RNA (gRNA) and a catalytically inactive form of Cas9 (dCas9). The tagged locus is then affinity-purified to enable identification of associated molecules. Locus-tagging can be achieved in cells by expressing engineered DNA-binding molecules (Fujita *et al.* 2013; Fujita & Fujii 2013, 2014b; Fujita *et al.* 2015; Fujita *et al.* 2016a; Fujita *et al.* 2017b), or *in vitro* using recombinant or synthetic engineered DNA-binding molecules (Fujita & Fujii 2014a; Fujita *et al.* 2016b; Fujita *et al.* 2017a) Combination of enChIP with mass spectrometry (MS), RNA sequencing, and next-generation sequencing (NGS) enables identification of proteins (Fujita *et al.* 2013; Fujita & Fujii 2013, 2014b), RNAs (Fujita *et al.* 2015), and other genomic regions (Fujita *et al.* 2017a; Fujita *et al.* 2017b) that interact with specific loci of interest in a non-biased manner.

To perform locus-tagging in primary cells, it is necessary to express both dCas9 and gRNA by transduction or other methods. However, the low transduction efficiency of some cell lineages results in a low percentage of cells expressing both dCas9 and gRNA. To resolve this technical issue, we generated transgenic (Tg) mouse lines expressing 3xFLAG-tagged *Streptococcus pyogenes* dCas9 (3xFLAG-dCas9), either constitutively or inducibly. To facilitate their use in various experimental contexts, expression of the tagged dCas9 and/or a reporter green fluorescent protein (GFP) can be flexibly induced or abolished by Cre- or FLPe-mediated site-specific recombination events that delete expression-modulating cassettes. We anticipate that these novel Tg mouse lines will serve as a powerful tool for efficient enChIP analysis in primary cells.

## Results and Discussion

### Generation of Tg mouse lines expressing 3xFLAG-dCas9

To facilitate enChIP analysis using primary mouse cells, we generated two Tg mouse lines expressing 3xFLAG-dCas9 (Fig. 1A, B). One line, 3xFLAG-dCas9-IRES-EGFP, harbors 3xFLAG-dCas9 and IRES-EGFP in the *Rosa26* locus (Fig. 1A). In the other line, 3xFLAG-dCas9 and IRES-EGFP are present at the *Rosa26* locus, but expression of 3xFLAG-dCas9 can be induced by Cre-mediated deletion of the STOP cassette (along with the *neo*^*r*^ gene), and EGFP expression can be disrupted by FRT-mediated deletion of the IRES-EGFP cassette (Fig. 1B (i)). 3xFLAG-dCas9/CTV (neo^+^/ GFP^+^) mice were crossed with CAG-Cre mice (Matsumura *et al.* 2004) to delete the STOP cassette and *neo*^*r*^ gene, yielding 3xFLAG-dCas9/CTV (GFP^+^) mice (Fig. 1B (ii)). 3xFLAG-dCas9/CTV (GFP^+^) mice can be further crossed with CAG-FLPe mice (Schaft *et al.* 2001) to delete the IRES-EGFP cassette, yielding 3xFLAG-dCas9/CTV mice (Fig. 1B (iii)). Targeted integration of transgenes was confirmed by PCR (Fig. 1C). All mice were viable and fertile with normal litter sizes and did not exhibit any morphological abnormalities. Expression of EGFP was observed throughout the body, including thymocytes and splenocytes (Fig. 2A), and 3xFLAG-dCas9 was detected in nuclear extracts (NE) of thymocytes (Fig. 2B). In conventional enChIP using primary cells from mice, it is necessary to transduce both tagged dCas9 and gRNA. To compare the number of Tg mice required for enChIP with that required for conventional enChIP, we isolated CD4^+^ T cells from a wild-type C57BL/6 mouse and transduced them with a retroviral plasmid expressing 3xFLAG-dCas9 (3xFLAG-dCas9/MSCV-EGFP). As shown in Supplementary Figure S1, the transduction efficiency of 3xFLAG-dCas9/MSCV-EGFP was about 10%. Considering that all cells express 3xFLAG-dCas9 in our Tg mice (Fig. 2A), this means that 10 times more mice are required for conventional enChIP than for enChIP using our Tg mice. Thus, our Tg mouse lines have the advantage of reducing the number of mice required for enChIP, as well the time and effort needed to perform enChIP analysis in primary mouse cells.

**Fig. 1.**
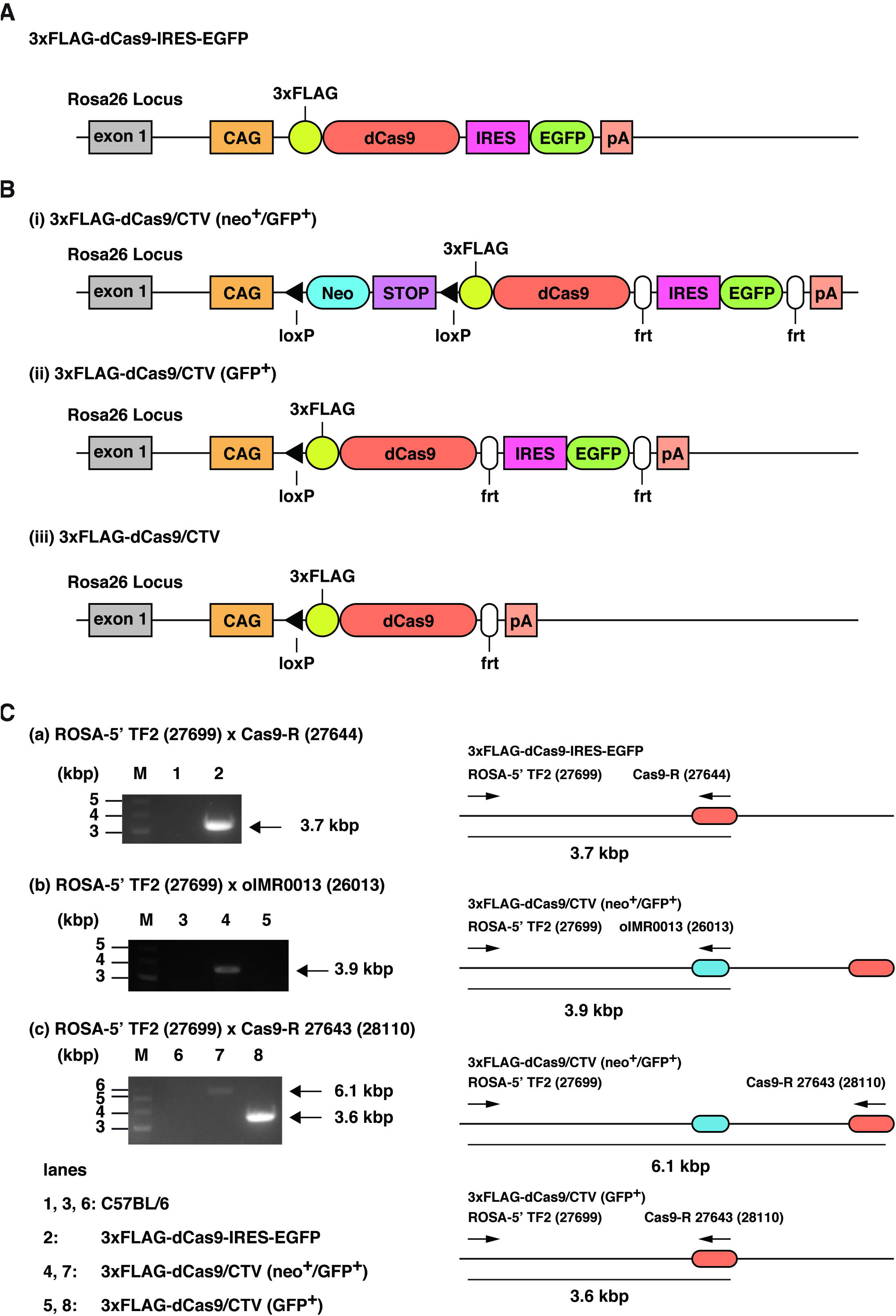
Generation of Tg mouse lines expressing 3xFLAG-dCas9. (**A**) Scheme of the targeted locus of 3xFLAG-dCas9-IRES-EGFP. (**B**) Scheme of the targeted loci of (i) 3xFLAG-dCas9/CTV (neo^+^/GFP^+^); (ii) 3xFLAG-dCas9/CTV (GFP^+^); and (iii) 3xFLAG-dCas9/CTV. (**C**) Genotyping PCR of Tg mouse lines.

**Fig. 2.**
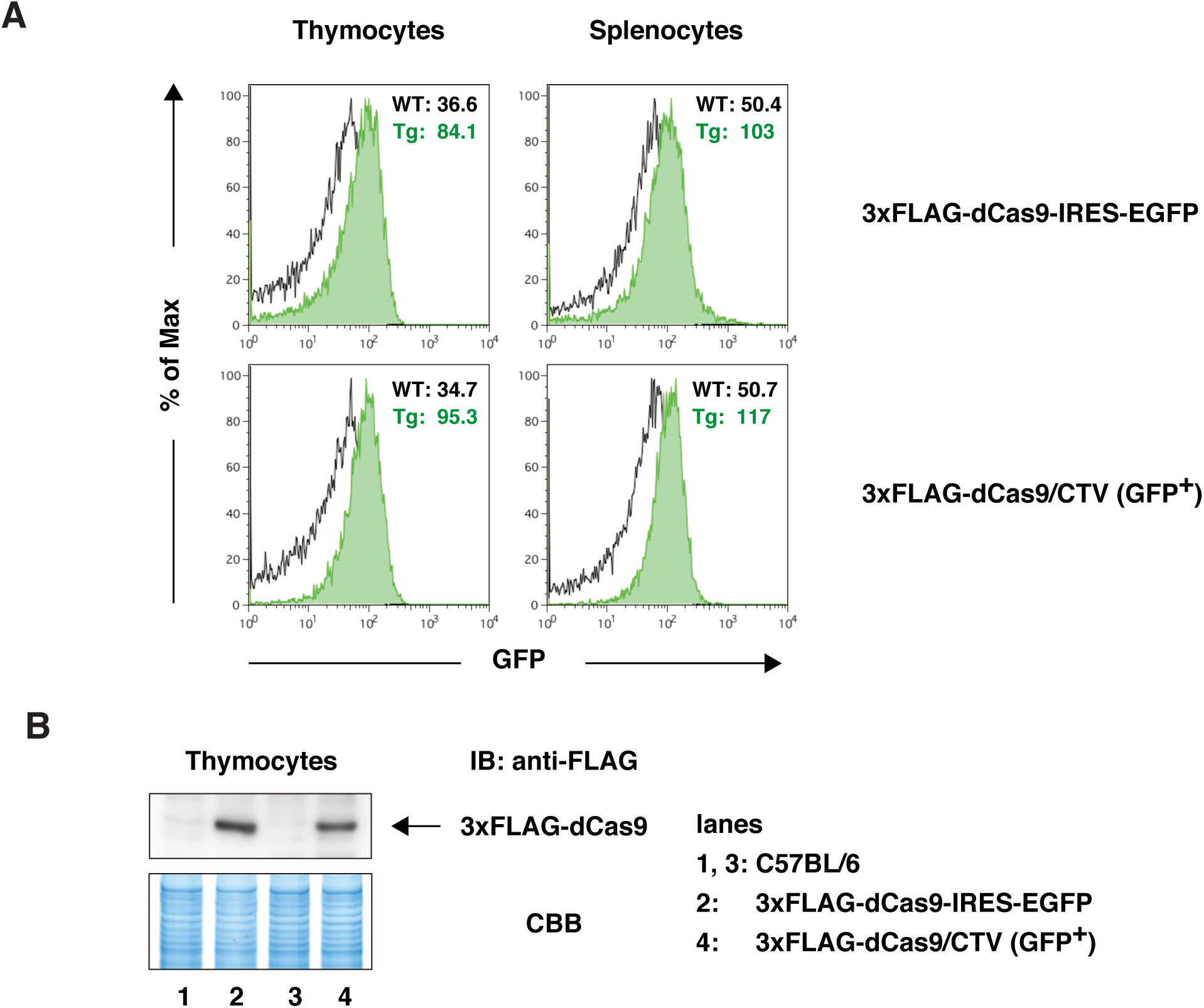
Expression of 3xFLAG-dCas9. (**A**) Expression of GFP in thymocytes and splenocytes from Tg mice. Fluorescence in the FL-1 channel (GFP) is shown for C57BL/6 mice (WT: black) and Tg mice (Tg: green). Numbers represent mean fluorescence intensities (MFI) in the FL-1 channel. (**B**) Expression of 3xFLAG-dCas9 in thymocytes from Tg mice. Expression of 3xFLAG-dCas9 was detected by immunoblot analysis with anti-FLAG Ab. Coomassie Brilliant Blue (CBB) staining is shown as a protein loading control.

### enChIP analysis using primary CD4^+^ T cells

Next, we performed enChIP analysis using primary cells from the Tg mice (Fig. 3A). For these experiments, CD4^+^ T cells were purified from 3xFLAG-dCas9-IRES-EGFP mice and 3xFLAG-dCas9/CTV (GFP^+^) mice and activated with anti-CD3 and anti-CD28 Abs. The activated cells were transduced with a retroviral vector expressing gRNA targeting the c-*myc* promoter (m-c-*myc* gRNA #1/pSIR-hCD2) or a negative control vector (pSIR-hCD2), and 2 days later, hCD2^+^ cells were isolated and expanded in media containing IL-2. Cells were fixed with formaldehyde and subjected to enChIP analysis using anti-FLAG Ab. Yields of enChIP were monitored by real-time PCR. As shown in Figure 3B and C, efficient enrichment of the c-*myc* promoter region, but not irrelevant loci (*Gapdh, Pax5*), was detected in samples expressing gRNA targeting the c-*myc* locus. By contrast, no enrichment was observed for samples in the absence of gRNA. The yields of enChIP were comparable between 3xFLAG-dCas9-IRES-EGFP mice and 3xFLAG-dCas9/CTV (GFP^+^) mice. These results demonstrate that primary cells from these Tg mice can be used for enChIP analysis.

**Fig. 3.**
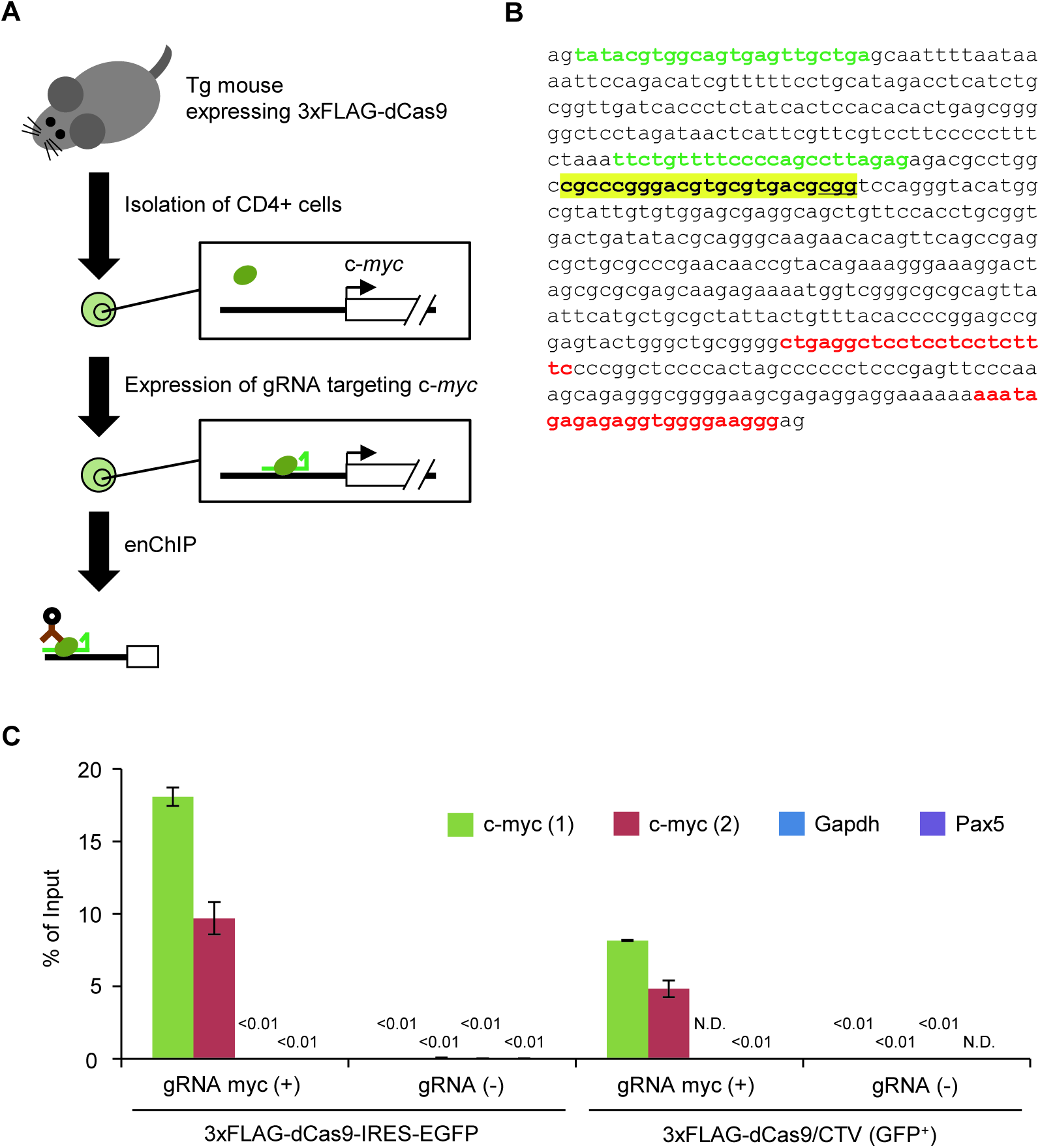
enChIP analysis of CD4^+^ T cells from 3xFLAG-dCas9 Tg mice. (**A**) Scheme of enChIP analysis using Tg mouse lines. (**B**) Positions of primer sets used for enChIP real-time PCR. Green letters: primer positions for c-myc (1); red letters: primer positions for c-myc (2); yellow highlight: gRNA target site; underline: PAM. (**C**) Yields of enChIP analysis for the target site. Error bars represent differences between duplicate analyses. N.D.: not detected. Irrelevant loci (*Gapdh* and *Pax5*) were analysed as negative control loci.

### Conclusions

We generated Tg mouse lines expressing 3xFLAG-tagged dCas9 and retrovirally transduced gRNA targeting a genomic locus into primary CD4^+^ T cells from these mice. Using this approach, high yields of enChIP could be achieved. Thus, these Tg mouse lines represent a useful tool for enChIP analysis in primary mouse cells. The injection of adenovirus-mediated gRNA (Platt *et al.* 2014) into these Tg mice should also enable the isolation of genomic regions of interest from mouse tissues without the need for primary cell cultures. In addition, the Tg mouse strains generated in this study could be used for CRISPR interference (CRISPRi) experiments (Qi *et al.* 2013) using primary mouse cells. However, in enChIP analysis, such CRISPRi effects might be problematic. Therefore, it would be better to choose gRNA target sequences that bind to the CRSIPR complex without interfering with the functions of the target genomic regions.

## Experimental procedures

### Plasmids

To construct pCAG1.2-PM, a modified pCAGGS plasmid (Niwa *et al.* 1991) was digested with *Sac*I. Two oligonucleotides, *Mlu*I–*Pme*I oligo-S (27551) and *Mlu*I–*Pme*I oligo-A (27552), were annealed, phosphorylated, and inserted into the digested plasmid, yielding two plasmids, pCAG1.2-PM (*Pme*I–*Mlu*I) and pCAG1.2-MP (*Mlu*I–*Pme*I), distinguished by the orientations of the oligonucleotides. To construct 3xFLAG-dCas9/pCAG1.2-PM, pCAG1.2-PM was digested with *Eco*RV and *Not*I. 3xFLAG-dCas9/pMXs-puro (Addgene #51240) was digested with *Pac*I, blunted, and further digested with *Not*I. The vector backbone and the coding sequence of 3xFLAG-dCas9 were purified by agarose gel electrophoresis and ligated. To construct 3xFLAG-dCas9-IG/pCAG1.2-PM, 3xFLAG-dCas9/pCAG1.2-PM and 3xFLAG-dCas9/pMXs-IG (Addgene #51258) were digested with *Not*I and *Sal*I, respectively. After blunting, the plasmids were further digested with *Asc*I. The vector backbone and the coding sequence of 3xFLAG-dCas9 were purified by agarose gel electrophoresis and ligated. To construct 3xFLAG-dCas9-IG/pSKII-ROSA, pSKII-ROSA26arm0.5kb-zeo was digested with *Bam*HI and *Sal*I, and 3xFLAG-dCas9-IG/pCAG1.2-PM was digested with *Mlu*I and *Pac*I. After blunting, the vector backbone and the coding sequence of 3xFLAG-dCas9 were purified by agarose gel electrophoresis and ligated.

To construct 3xFLAG-dCas9/CTV, CTV (Addgene #15912) was digested with *Asc*I, and 3xFLAG-dCas9/pMXs-puro was digested with *Pac*I and *Not*I. After blunting, the vector backbone and the coding sequence of 3xFLAG-dCas9 were purified by agarose gel electrophoresis and ligated.

To construct 3xFLAG-dCas9/MSCV-EGFP (Addgene #82613), the MSCV-EGFP plasmid (DeKoter *et al.* 1998) was digested with *Hpa* I and ligated with the coding sequence of 3xFLAG-dCas9, which was derived from 3xFLAG-dCas9/pMXs-puro by digesting with *Pac* I and *Not* I followed by blunting with DNA Blunting Kit (Takara).

To construct a gBlock targeting the mouse c-*myc* promoter, the oligonucleotides mcMYC promoter 1 sense (27822) and mcMYC promoter 1 antisense (27823) were annealed, phosphorylated, and inserted into gRNA cloning vector *Bbs*I ver. 1 digested with *Bbs*I. To construct m-c-*myc* gRNA #1/pSIR, the gBlock was excised with *Xho*I and *Hin*dIII and inserted into *Xho*I/*Hin*dIII-cleaved pSIR (Clontech). To construct a retroviral vector for expression of gRNA against the mouse c-*myc* promoter (m-c-*myc* gRNA #1/pSIR-hCD2), pSIR-hCD2 (Addgene #51143) and m-c-*myc* gRNA #1/pSIR were digested with *San*DI and *Hin*dIII. The vector backbone and insert were purified by agarose gel electrophoresis and ligated.

Oligonucleotides used for construction of plasmids are shown in Table 1.

**Table 1.**
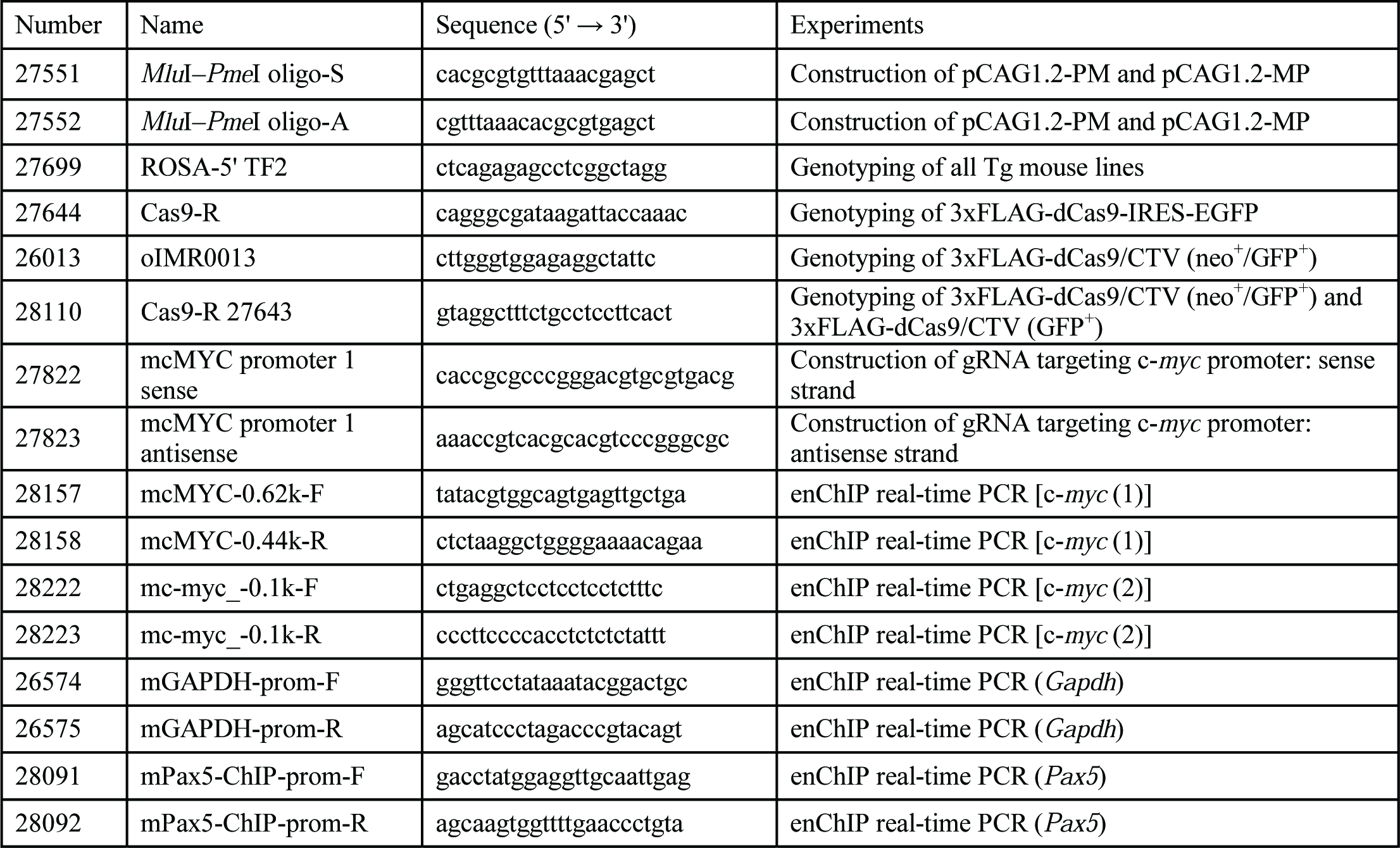
Oligodeoxyribonucleotides used in this study

### Mice

Embryonic stem (ES) cells (EGR-101) (Fujihara *et al.* 2013) were transfected with linearized 3xFLAG-dCas9-IG/pSKII-ROSA, as described previously (Oji *et al.* 2016). ES cells retaining the transgene in the *Rosa26* locus were injected into blastocysts (ICR × ICR) to generate chimeric mice. The chimeric mice were crossed with B6D2F1 mice to generate heterozygous 3xFLAG-dCas9-IRES-EGFP mice (strain name: B6D2-*Gt(ROSA)26Sor*^*em1(CAG-3XFLAG-dCas9,-EGFP)Osb*^) (RIKEN BioResource Center RBRC09976; Kumamoto University Center for Animal Resources and Development (CARD): CARD ID 2531).

B6JN/1 ES cells (Moriyama *et al.* 2014), derived from a hybrid between C57BL/6J and C57BL/6N, were electroporated with 3xFLAG-dCas9/CTV plasmid linearized with *Asi*SI. To generate chimeric mice, ES cells containing the *neo*^*r*^ gene, transgene, and *GFP* gene in the *Rosa26* locus were injected into blastocysts (Balb/c × Balb/c) by the animal facility group at RIKEN IMS. The chimeric mice were crossed with C57BL/6 mice to generate heterozygous 3xFLAG-dCas9/CTV (neo^+^/GFP^+^) mice (strain name: C57BL/6-*Gt(ROSA)26Sor*^*tm1(CAG-3XFLAG-dCas9,-EGFP)Hfuj*^). The 3xFLAG-dCas9/CTV (neo^+^/GFP^+^) mice were crossed with CAG-Cre mice (Matsumura *et al.* 2004) to generate 3xFLAG-dCas9/CTV (GFP^+^) mice (strain name: C57BL/6-*Gt(ROSA)26Sor*^*tm1.1(CAG-3XFLAG-dCas9,-EGFP)Hfuj*^). The 3xFLAG-dCas9/CTV (GFP^+^) mice can be crossed with CAG-FLPe mice (Schaft *et al.* 2001) to generate 3xFLAG-dCas9/CTV mice (strain name: C57BL/6-*Gt(ROSA)26Sor*^*tm1.2(CAG-3XFLAG-dCas9)Hfuj*^).

All animal experiments were approved by the Institutional Animal Care and Use Committee at Research Institute for Microbial Diseases, Osaka University.

### Genotyping

For genotyping, genomic DNA was extracted and subjected to PCR with KOD FX (Toyobo). PCR conditions were as follows: heating at 94°C for 2 min; followed by 38 cycles of 98°C for 10 s, 62°C for 30 s, and 68°C for 4 min. Primers used for genotyping PCR are shown in Table 1.

### Cell staining and flow cytometry

Thymi and spleens were isolated from euthanized mice and used to prepare single cells. For surface staining, cells were stained for 30 minutes at 4°C with fluorochrome-conjugated antibodies (Abs): fluorescein isothiocyanate (FITC)-conjugated mouse CD4 (130-102-541, Miltenyi) and phycoerythrin (PE)-conjugated human CD2 (hCD2) (347597, BD Bioscience). Flow cytometric analysis was performed on a FACSCalibur (BD Biosciences) and analyzed with the FlowJo software (TreeStar).

### Immunoblot analysis

NE were prepared with NE-PER Nuclear and Cytoplasmic Extraction Reagents (Thermo Fisher Scientific). Aliquots of NE (10 μg) were subjected to immunoblot analysis with anti-FLAG M2 Ab (F1804, Sigma-Aldrich), as described previously (Fujita & Fujii 2013).

### Transduction of gRNA and isolation of transduced cells

Transduction of retroviral expression plasmids into primary CD4^+^ T cells was performed as described previously (Naoe *et al.* 2007). Briefly, CD4^+^ T cells were purified from spleens using the Mouse CD4^+^ T cell isolation kit (Miltenyi, 130-104-454). Purified CD4^+^ T cells were activated with anti-CD3 Ab (3 μg/ml, 145-2C11, 553057, BD Pharmingen) and anti-CD28 Ab (3 μg/ml, 37.51, 553295, BD Pharmingen). A retroviral expression plasmid was transfected into Plat-E cells (Morita *et al.* 2000) along with pPAM3 (Miller & Buttimore 1986) using Lipofectamine 3000 (Invitrogen) to produce retroviral particles. Activated CD4^+^ T cells were transduced with the retroviral particles by the spin infection method (Naoe *et al.* 2007). After culturing for 2 days in RPMI complete media containing mouse IL-2 (20 ng/ml, 402-ML, R & D Systems), transduced cells were analyzed by flow cytometry. hCD2^+^ cells were purified using human CD2 MicroBeads (130-091-114, Miltenyi) and used for enChIP analysis.

### enChIP real-time PCR

enChIP real-time PCR was performed as described previously (Fujita & Fujii 2013) with some modifications. Briefly, the CD4^+^ T cells (ca. 1 × 10^6^) were crosslinked with 0.1% formaldehyde in RPMI complete media at 37°C for 10 min. After quenching and washing with PBS, the chromatin fraction was extracted and fragmented by sonication. The sonicated chromatin was used for enChIP using 2 μg of anti-FLAG M2 Ab. DNA was purified using ChIP DNA Clean & Concentrator (Zymo Research) and subjected to real-time PCR. Primers used in the analysis are shown in Table 1.

## Acknowledgments

We thank S. Muroi for genotyping ES cells, T. Ishikura for injection of ES cells into blastocysts, and K. Rajewsky and H. Singh for providing plasmids (Addgene plasmid # 15912 and MSCV-EGFP, respectively).

## Funding

This work was supported by the Takeda Science Foundation (T.F.), Grant-in-Aid for Scientific Research (C) (#15K06895) (T.F.), and Grant-in-Aid for Scientific Research (B) (#15H04329) (T.F., H.F.), ‘Transcription Cycle’ (#15H01354) (H.F.) from the Ministry of Education, Culture, Sports, Science and Technology of Japan.

## Abbreviations

enChIP: engineered DNA-binding molecule-mediated chromatin immunoprecipitation
CRISPR: clustered regularly interspaced short palindromic repeats
dCas9: catalytically inactive form of Cas9
gRNA: guide RNA
GFP: green fluorescent protein
MS: mass spectrometry
NGS: next-generation sequencing
Tg: transgenic

## Conflicts of interests

T.F. and H.F. have patents on enChIP (“Method for isolating specific genomic region using molecule binding specifically to endogenous DNA sequence”; patent number: Japan 5,954,808; patent application number: WO2014/125668). T.F. and H.F. are founders of Epigeneron, Inc.

## Availability of data and materials

All data generated or analyzed during this study are included in the published article. Tg mice generated in this study can be obtained from RIKEN BioResource Center and Kumamoto University Center for Animal Resources and Development (CARD).

## Authors’ contributions

H.F. designed and performed experiments (design and construction of transgenes, flow cytometric analysis, immunoblot analysis, transduction of retroviruses), wrote the manuscript, and directed and supervised the study. T.F. and M.Y. performed enChIP analyses. N.T. constructed the retrovirus vector expressing gRNA targeting the c-*myc* locus, and performed transduction of gRNA retroviruses and enChIP analysis. F.K. and M.Y. maintained the mouse colony. F.K. screened ES cells and genotyped mice. A.O. and M.I. performed CRISPR-mediated knock-in of 3xFLAG-dCas9-IRES-EGFP transgenes. I.T. generated ES cells harboring the 3xFLAG-dCas9/CTV transgene, chimeric mice, and knock-in mice.

